# Accelerating Protein Structure Prediction using Particle Swarm Optimization on GPU

**DOI:** 10.1101/022434

**Authors:** Hamed Khakzad, Yasaman Karami, Seyed Shahriar Arab

**Affiliations:** Computer Engineering Department, Iran University of Science and Technology, Tehran, IRAN; Department of Biophysics, Faculty of Biological Sciences, Tarbiat Modares University, Tehran, IRAN

**Keywords:** Protein structure prediction, Graphical processing unit, Particle swarm optimization, Ab-initio, PSI-Blast

## Abstract

Protein tertiary structure prediction (PSP) is one of the most challenging problems in bioinformatics. Different methods have been introduced to solve this problem so far, but PSP is computationally intensive and belongs to the NP-hard class. One of the best solutions to accelerate PSP is the use of a massively parallel processing architecture, such graphical processing unit (GPU), which is used to parallelize computational algorithms. In this paper, we have proposed a parallel architecture to accelerate PSP. A bio-inspired method, particle swarm optimization (PSO) has been used as the optimization method to solve PSP. We have also performed a comprehensive study on implementing different topologies of PSO on GPU to consider the acceleration rate. Our solution belongs to ab-initio category which is based on the dihedral angles and calculates the energy-levels to predict the tertiary structure. Indeed, we have studied the search space of a protein to find the best pair of angles that gives the minimum free energy. A profile-level knowledge-based force field based on PSI-BLAST multiple sequence alignment has been applied as a fitness function to calculate the energy values. Different topologies and variations of PSO are considered here and the experimental results show that the speedup gain using GPU is about 34 times faster than CPU implementation of the algorithm with an acceptable precision. The energy values of predicted structures confirm the robustness of the algorithm.

## 1. Introduction

Bioinformatics is the science of using computer and information technology in biology. One of the main challenges in structural bioinformatics is the prediction of protein tertiary structure which aims to find the 3D structure with the lowest free energy of a protein from its sequence. The significance of PSP is that scientists believe the function of a protein in a cell depends on its folding and structure [1]. Findings of the native conformation can be used in designing drugs, recognizing the function of inheritance and contagious diseases and prevention of some diseases like Alzheimer, cystic fibrosis and mad cow [2]. However, common computational methods are unable to solve PSP problem as their computational time and complexity are both expensive. At the other hand, experimental methods for PSP based on NMR and X-ray despite of significant development [3] are still time consuming and expensive. The computational methods are divided into three groups [4]: homology modeling, threading, and ab-initio. The first two are knowledge-based methods using the information of known protein structures, but the third is based on physicochemical parameters and solves the problem by inspiring what is happening in the nature; they predict the 3D structure of proteins given their primary sequences.

GPU was first introduced for graphical purposes. However, as its architecture is based on Single-Instruction Multiple-Data (SIMD) so data-parallel algorithms could be implemented on these devices to get acceptable speedup. In order to perform general-purpose computing on GPU, CUDA has been developed which has greatly simplified programming on GPU. Recently, GPU has already been used to accelerate plenty of application in bioinformatics, such as CUDASW++ [5], GHOSTM [6], and MetaBinG [7]. In addition, an accelerated GPU-based version of the well-known NCBI-BLAST tool has been developed which achieves 3-4 times speedups [8]. The CUDA-BLASTP has also been developed which is 10 times faster than the CPU implementation [9].

PSO is a population-based stochastic algorithm for optimization of continuous nonlinear functions. It is based on social behavior of birds or fishes. Unlike evolutionary algorithms, PSO does not use selection. Indeed, all population members survive from the beginning of a trial until the end. Their interactions result in iterative improvement of the quality of problem solutions over time [10]. PSO is simpler than standard GA, producing results in shorter time with less computation cost [11]. PSO has also been significantly applied in a wide range of problems in computational biology and bioinformatics especially on protein structure prediction [12 13 14].

There are significant implementations of PSO on GPUs. Li et al. [15] proposed a Fine-Grained parallel PSO method based on GPU acceleration. This method eliminates standard FGPSO drawbacks by increasing the population size and speeds up its execution by using GPU as an easy accessible device. Zhou et al. [16] also parallelized Standard PSO using GPU. The proposed GPU-SPSO method achieved at most 11 times speed up in comparison to CPU implementation on four classical benchmark test functions when the swarm population size was 20000. Furthermore, Souza et al. [17] implemented PSO on GPU and obtained about 3 times speed up on two different test functions.

In this paper, Our solution is based on Anfinsen thermodynamic hypothesis that claims the native structure of a protein is the conformation that has the minimum free energy in the physiological environment [18]. Based on this hypothesis, PSP belongs to the category of optimization problems with huge number of conformations in solution space. In literature, different representations for the structure of proteins are introduced to reduce the complexity of the problem based on different energy equations. One simple model is the Hydrophobic-Polar (HP) lattice model that considers the hydrophobicity feature of amino acids and divides them into two groups of H’s (Hydrophobic) and P’s (Polar). This model has been used widely due to its simplicity, but the results are not exact and accurate [19 20]. Another common and simple model is AB-off lattice which is also based on hydrophobicity. The energy calculations in this model are based on the hydrophobicity, bend angles and the distance between amino acids which makes it more accurate [21 22]. Other models use the detailed representation of proteins with all the corresponding atoms, based on force fields such as CHARMM, AMBER, GROMOS and etc. Meanwhile, knowledge-based potentials are quite successful in fold recognition and ab-initio structure prediction [23]. These force fields are fast and simpler and are derived from known protein structures using statistical analysis. Force fields could calculate the energy of a protein based on atom-level [24 25] or residue-level [26 27] interactions. Atom-level force fields are more exact though being more complex. However, another group is introduced which is based on profile-level knowledge-based potentials. This group is more accurate than residue-levels and has reduced complexity compared with atom-levels [28].

In our approach, we have studied the search space of a protein strongly to find the best pair of dihedral angles that gives the minimum free energy. A profile-level knowledge-based force field has been applied to energy calculation as the fitness function. PSI-blast is used to construct the profiles from multiple sequence alignments. Then, the knowledge-based force field is constructed from these profiles to measure different types of potentials for any proteins. In fact, this ab-initio method is based on the dihedral angles and calculates the energy-levels to predict the tertiary structure. The crucial objective in this paper is the acceleration of PSP by using GPU which yields about 34 times speedups.

## 2. Material and Methods

### a. Force Field

To calculate energy values, the profile-level knowledge-based Φ/Ψ dihedral angle potential is used here. The profile of amino acids sequences are constructed based on multiple sequence alignment with PSI-BLAST against the NR90 database, and the frequency profiles are obtained from the output of PSI-BLAST. In this case, it is a matrix with 20 columns (one for each amino acid types) and the number of rows is equal to the sequence length. Every member of this matrix shows the occurrence probability of every amino acid in each position of the sequences. These frequency profiles are converted into binary profiles by a threshold. Then, binary profiles are used to calculate the dihedral potential. This potential is based on the propensity of each pair of torsion (Φ/Ψ) angels for each dihedral class and the total number of torsion angle bins is assumed to be 36. The dihedral potential is calculated as Equation 1:

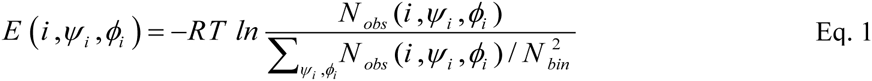

where i is the interaction type and Φ_i_, Ψ_i_ are the torsion angles. The torsion potential is the logarithm of the number of observed occurrences of the interaction center type i at torsion angles of (Φ_i_, Ψ_i_) normalized by the averaged occurrence. Finally, this calculation is used as fitness function in our PSO algorithm.

### b. PSO Algorithm

In comparison to the algorithms that are used to find the near optimal solutions, PSO produces acceptable results. PSO concept is close to evolutionary algorithms whilst requires simpler programming to be implemented. Every particle of the PSO is one solution of the search space. The aim of this algorithm is that the whole particle reaches the optimal global point. In this way, the speed of each particle changes dynamically, considering the last move and the neighbor particles. Therefore, position and velocity vectors are X_i_ = (X_i1_,…,X_id_) and V_i_ = (V_i1_,…,V_id_) for the i^th^ particle in the D-dimensional search space, respectively. Velocity changes give each particle the ability to search around the best local or global point. Equation 2 and 3 indicate a common PSO (CPSO) algorithm to update the position and velocity recursively:

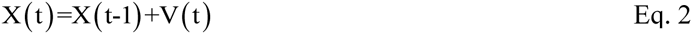

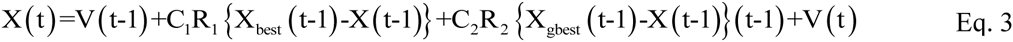

where V(t) and X(t) represent the velocity and the position of one particle at time t, respectively. X_best_ (t-1) is the local best position reached by the particle up to time t-1 and X_gbest_(t-1) is the global best point found by the whole swarm. C_1_ and C_2_ are two positive constants, while R_1 and_ R_2_ are two random numbers uniformly distributed between 0 and 1. From Equation 3, it is understood that the speed of each particle is updated based on its previous speed, its best position up to now (cognitive component) and the global optimal position (social component).

After the position update process, fitness function is calculated for each particle and the local best and global best values will be updated. Other variations of PSO are also introduced to improve the result and its efficiency [29]. Indeed, the “Inertia Weight” which is a parameter to adjust the impact of the velocity on the result is considered. So the velocity will be updated as Equation 4:

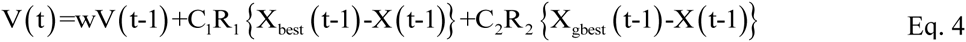

where “w” could be set as a constant or be changed based on different linear or nonlinear functions. Larger values of inertia weight help particles to search the whole space while lower values help better local search.

### c. CUDA Architecture

Compute Unified Device Architecture (CUDA) is a general-purpose parallel computing architecture introduced by NVidia in 2006, as a tool to GPU programming. Its architecture is based on multiple threads, called Single-Instruction Multiple-Thread (SIMT). Parallel programs use an extension of C language to get developed in CUDA. Most of the programs have a data-parallel portion that could be run in parallel and have also a sequential part. CPU has the responsibility to run the serial portion of the code, reading and writing the data files, preparing input data for GPU, accepting the result of GPU and storing answers and solutions. The data-parallel portion runs on the GPU and is assigned a function called kernel on CUDA that maps to a grid on the GPU. Every grid is composed of thread blocks that have specified number of threads. Kernels have instructions that are executed by all the threads of the specified grid in parallel, but different kernels in the entire algorithm are executed sequentially. The number of thread blocks and threads used by every kernel should be determined on the CPU when it’s called.

In CUDA memory model, each thread could access its own local memory and register. Also, all the threads in a thread block have access to that block’s shared memory. Constant and Texture memories are read only memories that could be accessed by all the threads of a grid and the large Global memory belongs to all the threads. GPUs based on their types, have some multi-processors, each of which has 8 processors or cores. Each multi-processor has 16Kb shared memory, 16Kb registers and 64Kb constant memory. Also, multi-processors could access the global device memory and texture memory for larger data storage [30]. The design and implementation of different algorithms on the GPU should be decided according to the limited amount of memories and their different performance.

### d. Proposed PSO-GPU Method

PSO-GPU is a search method based on parallel particles that move independently in the problem search space. Every particle is one possible structure for the protein and the position vectors are the vectors of dihedral angles (Phi and Psi). Velocity vectors show the rate of getting close to the answer and changing these angles. The algorithm starts with a first population generated by a novel pseudo-random method. The proposed method prevents the divergence of first generation from the native structure, while increases the convergence speed of the algorithm. In this method, a database of all dihedral angles of amino acids for about 9000 proteins with known 3D structures-from different families- is constructed. Considering triple combinations of residues in the sequence, twenty databases will be made from the first one each of which is related to one amino acid. Every database includes 400 different sets of angles for all the possible combinations of three amino acids in a way that the first one is fixed and the other two are changed. To construct the first generation based on this method, one should first determine the relevant database for every amino acid and its two following amino acids, then one pair of φ and Ψ angles should be selected randomly. As a result, the complete random selection of angles is turned into a selection that is based on previous experimental observations. This part causes no overhead due to the fact that it is used only once at the beginning of the algorithm.

After that, the energy values of each particle will be evaluated using a knowledge-based potential force field. Then, the iterative process begins as follows: (1) Position and velocity vectors are updated recursively using Equation 2 and 3 or 4. (2) The energy values of every particle are calculated. (3) Local and global best positions are updated. This process continues until the best conformation that has the lowest energy value will be generated. This conformation has the most appropriate dihedral angles, close to the native structure for the protein being predicted. The restriction considered here for the dihedral angles is their degree of freedom that should be among -180 to +180. In order to consider this condition, we set two different equations to check the boundaries. Equation 5 shows Fixed Boundary (FB) check and Equation 6 shoes Periodic Boundary (PB) check:

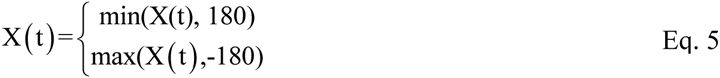

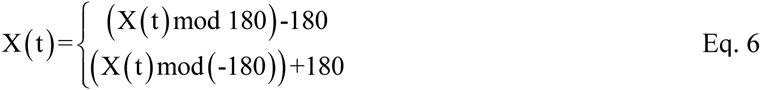

To implement this algorithm on the GPU, different kernels are determined for every part of the algorithm. The problem dimension here is equal twice the sequence length because each dimension shows one of the two dihedral angels for every residue. First population that is generated on the CPU is copied from CPU, in addition to the knowledge-based force field, to the global memory of GPU. Considering Ring PSO [31] each particle is assigned one block. Based on this architecture, the number of blocks used for the position update kernel is equal to the number of conformations and the number of threads per block is twice the sequence length. Also, coalescing is considered here; position and velocity vectors could be accessed with half-warp threads. The number of blocks used for the fitness evaluations is also equal to the number of conformations and number of threads per block is equal to the sequence length because the energy calculations could run in parallel for every residue just by reading the corresponding dihedral angels of that residue from the global memory of position vectors.

At the end, the final global best position vector determines the best conformation for the protein to be predicted. Because data transfers occur only at the beginning and at the end of the algorithm, remarkable speedup is achieved. Also, to reduce the amount of used global memory, only the necessary parts of the force field are transferred to the GPU. As the sequence of the protein to be predicted is fixed during the prediction process, it could be determined that which parts of the force filed will be used based on the binary profile of each residue. The only part in this algorithm that needs particles to share data is the function that updates the global best positions. To parallelize this part, different topologies are introduced. In this paper, Ring, Star and Global topologies are implemented to compare the result. Figure 1 shows these topologies. Every particle possesses a vector that shows its local best position. At the end of each iteration, one kernel compares the current fitness value of each particle, with the fitness value of its best position to determine the local best for that particle.

**Figure 1.**
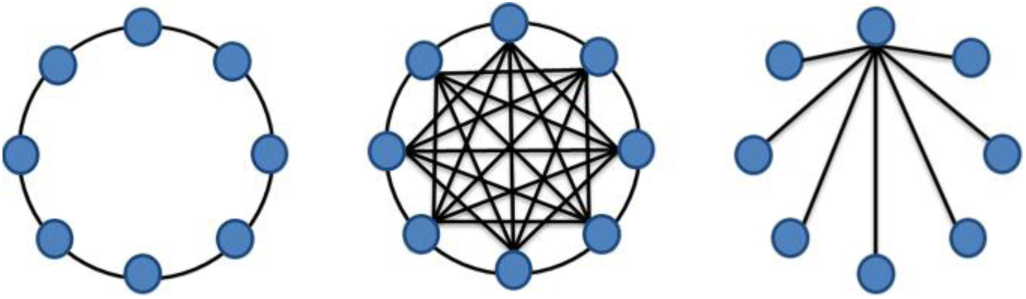
Ring, Global, and Star topologies of PSO.

In the Ring topology, the kernel that corresponds to updating the local best position, also updates the global best position for every particle. One block is assigned to this kernel and each thread of the block reads three fitness values of its corresponding particle’s conformation and the two neighbors on the right and left to determine the best global particle. In the star topology, one particle is responsible to update the global best value in each iteration. This particle receives the current fitness values from every particle and compares these values to find the best of them. In the global topology, every particle is considered to search the whole swarm and find the global best fitness value. Based on global topology, results are converged faster to the native structure. At the same time because of its fully connected structure, there is more communication between particles which takes more time to produce the result.

## 3. Results and Discussions

### a. Case Study

The PSO-GPU algorithm is implemented on a PC with a Quad-core AMD Opteron Processors (2.1 GHz), a Tesla C1060 with 240 thread processors and 3.12 GB of RAM. Parallel implementations are done under CUDA 4 and are compared with the sequential one, on the same PC. The PSO parameters are set as follows: w=0.729844 and C_1_=C_2_=1.49618. Different protein structures with different lengths from Protein Data Bank (PDB) are tested as follows: 1CRN (46), 1A1X (106), 3NPD (111) and 3RFN (151). We’ve implemented Common PSO (CPSO) and constant Inertia weight PSO (IPSO). Also, for both of these variations we considered Fixed Boundary (FB) check and Periodic Boundary (PB) check on the GPU to bring better comparison. Although the Tesla C1060 has the ability to use double precision arithmetic to increase numerical reproducibility, we consider the single precision due to its faster performance [32]. However, the variance of outputs in 10 independent runs was not notable. Finally, the execution time is averaged over 10 independent runs and the best fitness values are selected for each parallel or sequential implementation. Figure 2 shows the comparison between energy values of those PSO variations with different iteration numbers. Each part of this figure refers to one of the mentioned proteins.

**Figure 2.**
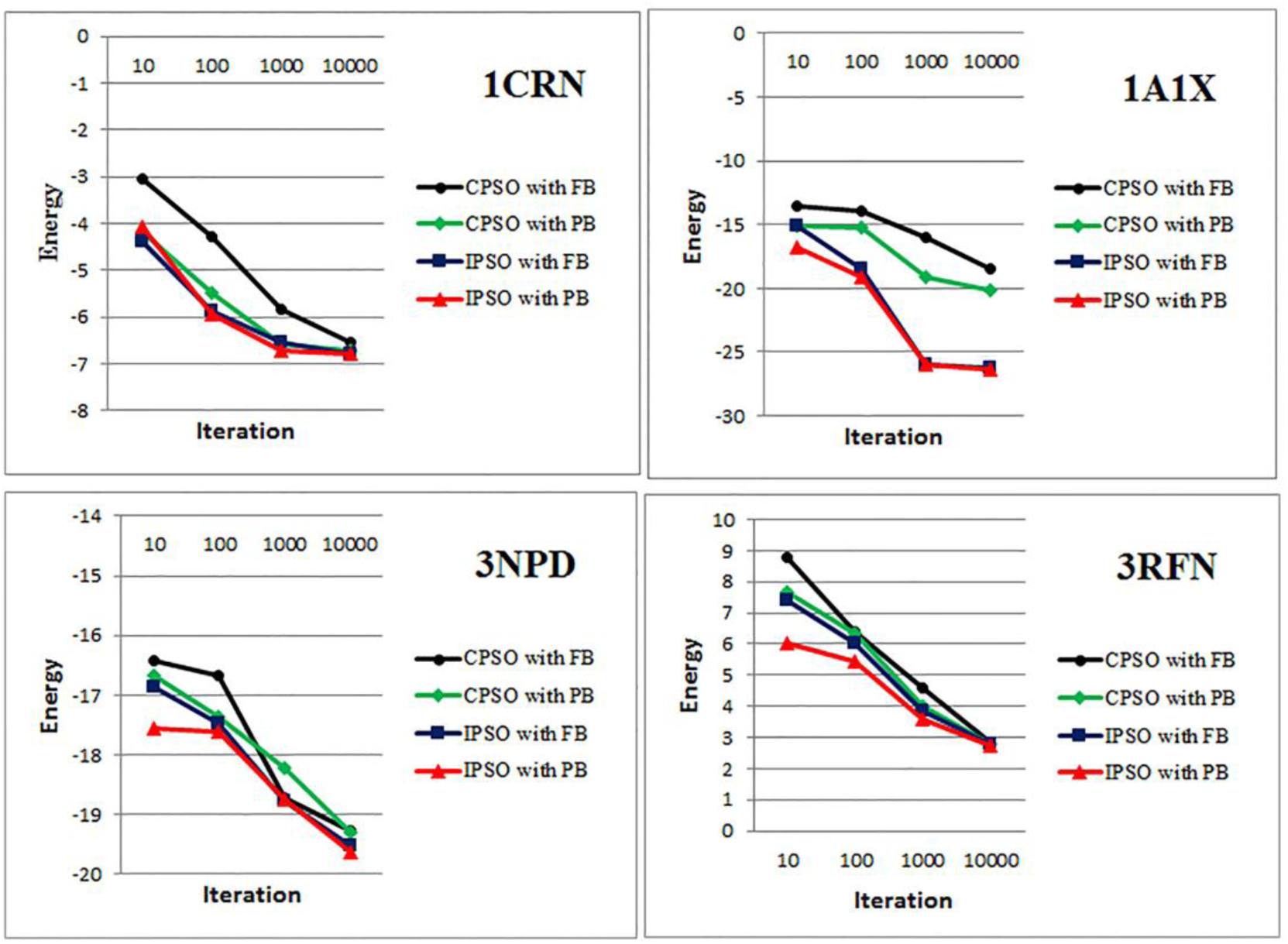
GPU-CPSO and GPU-IPSO implementation on proteins. In CPSO, the velocity vector is updated using Equation 3 without considering inertia weight. In IPSO, the w=0.729844 (according to the experiments) is defined. The results are shown according to the algorithm iteration. Accordingly, IPSO shows better convergence to the native energy in comparison with CPSO. The periodic boundary check parameter is also works better than fixed boundary check, respectively.

The comparison between CPU and GPU time for parallel IPSO with PB check is shown in Table 1. The number of considered iterations is 10000 with 512 particles (protein conformations) in this implementation. In fact, the number of particles and the sequence length is limited to the GPU size; in the case of Tesla C1060, the upper bound for the number of particles is 512. Problem dimension is twice the sequence length because it includes dihedral angles of phi and psi. As it is shown in the table, the GPU implementation of the algorithm is about 34 times (average) faster than the sequential implementation on CPU.

**Table 1.** Speedups achievement using CPU-GPU time comparison.

| Protein | Sequence Length | Native Energy | Predicted Energy | CPU Time(s) | GPU Time(s) | Speedup |
| --- | --- | --- | --- | --- | --- | --- |
| 1CRN | 46 | -6.8342 | -6.8 | 150 | 4 | 36 |
| 1A1X | 106 | -26.4986 | -26.31 | 297 | 9 | 32 |
| 3NPD | 111 | -19.6548 | -19.63 | 310 | 9 | 33.6 |
| 3RFN | 151 | 2.73666 | 2.73 | 455 | 12 | 36 |

Another comparison is also done between the dihedral energy values of the predicted and native structures. Generated conformation for each protein is validated due to the closeness of its energy value with the energy of native conformation. The dihedral energy values are calculated using the profile-level knowledge-based force field. As Table 1 indicates, the predicted conformations have acceptable energy values in comparison to the native structures. In PSP problem, different conformations for a protein could have the same energy value because each residue may have different dihedral angles. So the problem of PSP is multimodal and every particle of the PSO with the same energy value can be represented by different angles and rotations.

### b. PSO Topological Comparison

We have implemented the Ring, Star and Global topologies on GPU. As it is expected, the accuracy of the Global topology is better. As shown in figure 3, speedups for these topologies are obtained from the comparison of their GPU and corresponding CPU implementations. As for the different number of iterations, the parallel Ring PSO proved to be faster than parallel Global and Star PSO; all these GPU implementations with 10000 iteration were about 34 times faster than their corresponding CPU implementations.

**Figure 3.**
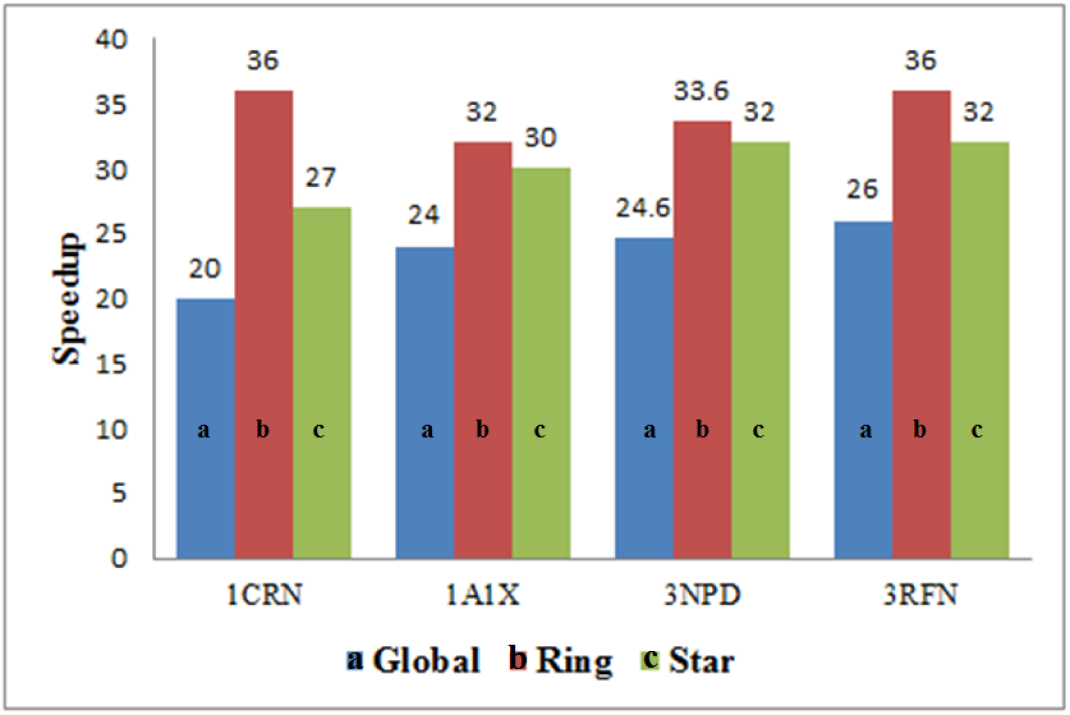
Speedup comparison between PSO topologies for different proteins with iteration = 10000.

### c. Comparison with Other Approaches

To the best of our knowledge, the combinatorial approaches based on PSO and GPU up to now, have not been applied for protein structure prediction. However, there are some methods which used PSO or GPU to tackle this problem separately. Therefore a general comparison is made through considering such approaches. Zhu et al. [22] proposed a parallel architecture for clonal selection algorithm on CUDA to predict the structure of proteins. In their work, two small Fibonacci sequences with length 13 and 21 were considered instead of real protein sequences. The speedup ratio was about 16. However, this work cannot fully verify the efficiency of their method because they did not test the algorithm on real protein sequence. In [33], parallel PSO was implemented in a distributed computing environment and the energy empirical function ECEPP/3 was used. It was tested on a known peptide, *leuenkephalin* with 24 torsion angles (5 amino acids) and another peptide called *[AAQAA]3Y* with 75 torsion angles (16 amino acids). Although, the proposed method was able to find the minimum energy, there was no comparison performed with the CPU implementation. Therefore, it is difficult to know whether this method overcome the computational complexity of PSP problem.

In [34] an evolutionary algorithm based on the standard harmony search strategy (PBHS) is parallelized using GPU to solve the PSP problem. However, this approach is implemented on AB-2D off lattice and just predicts the protein secondary structure to compare the standard harmony search with PBHS. Their results proved that the GPU implementation achieved about 68 x speeds up in comparison with CPU. Finally, Campeotto et al. [35] also proposed a multi-agents framework to solve PSP problem. They used concurrent agents on GPU to accelerate the prediction. Each agent has the ability to apply different kind of constraints to find the native structure. Three different kinds of constraints have been implemented in this work. The work is tested on 7 different proteins with sequence lengths of 12-100 amino acids. The average speed up they achieved is 26.93 which is less than the average speed up of our work (34.4).

## 4. Conclusion

The parallel implementation of one bioinformatics algorithm leads to the prediction of protein tertiary structure was introduced in this paper. PSO is one of the bio-inspired algorithms that have been used to predict the structure of proteins and produced acceptable results, but these algorithms are very slow when implemented on the CPU. GPU is a fast processing unit that is used to accelerate different computational algorithms these days. We have examined different proteins with different lengths to prove the correctness of our result. In this paper, we have considered two different boundary checks and compared CPSO with IPSO. In addition, three different Global, Star and Ring topologies have been implemented both on CPU and GPU. The results show that Ring PSO converges to the native structure faster, whereas the Global PSO is more accurate. Meanwhile, the Parallel PSO using CUDA runs about 34 times faster than the sequential implementation on the CPU. Also the predicted structure using IPSO with PB checks has energy values closer to the native.

